# Metabolic reprogramming regulates histone lactylation during zebrafish caudal fin regeneration

**DOI:** 10.1101/2024.09.28.615596

**Authors:** Jorge Borbinha, Raquel Lourenço, Ana S. Brandão, Ana Carvalho, Rune Matthiesen, António Jacinto

**Author notes:** Correspondence (A.J), (R.L.). These authors contributed equally to this work (Jorge Borbinha, Raquel Lourenço).

## Abstract

Tissue regeneration relies on precise molecular mechanisms controlling cell fate transitions, with cell metabolism emerging as a key regulator. Lactate-derived histone lactylation has recently been identified as an important epigenetic modification involved in the regulation of gene expression across various biological processes. In this study, we report an increase in global histone lactylation and H3K18Lac levels in the mesenchyme and osteoblasts of the zebrafish caudal fin, during the early stages of regeneration. Our findings demonstrate that this epigenetic modification is functionally regulated by increased lactate levels, while inhibition of glycolysis and lactate production significantly reduces histone lactylation. These results suggest that metabolic reprogramming, in response to caudal fin injury, regulates histone lactylation, potentially modulating gene expression essential for cell plasticity and proliferation. This study expands our understanding of how metabolic-epigenetic interactions influence regenerative processes, providing valuable insights for the development of novel therapeutic strategies to enhance tissue repair.

**Summary statement:** Lactate-derived histone lactylation plays a role in the early stages of zebrafish caudal fin regeneration, coupling metabolic adaptation with tissue repair mechanisms, through epigenetic regulation.

## Introduction

During tissue and organ regeneration there is a precise control over cell fate changes, like dedifferentiation to generate proliferative progenitors and redifferentiation to restore lost tissues. Recently, cell metabolism has emerged as a key driver of these transitions, with differentiated cells relying on oxidative phosphorylation for energy source, while stem and proliferative cells favour aerobic glycolysis with lactate production (Tatapudy et al., 2017). This change between metabolic states is described as metabolic reprogramming. Several reports, including our previous study, have shown that metabolic reprogramming plays a vital role in regenerative programmes (Osuma et al., 2018; Honkoop et al., 2019; Yucel et al., 2019; Varela-Rodríguez et al., 2020; Sinclair et al., 2021; Brandão et al., 2022). We identified a critical role for metabolism in triggering early events of zebrafish caudal fin regeneration, with bone-producing osteoblasts prioritizing glycolysis and lactate production during dedifferentiation (Brandão et al., 2022). Mature osteoblasts dedifferentiation begins as early as 6 hours post-amputation (hpa) alongside the initial wound healing response (Knopf et al., 2011; Sousa et al., 2011; Stewart & Stankunas, 2012). Osteoblasts re-enter the cell cycle and incorporate the blastema, a cluster of undifferentiated proliferating cells, by 12-24 hpa, and redifferentiate into mature osteoblasts during the outgrowth phase, from 48 hpa to 20 days post-amputation, completing fin regeneration (Stewart et al., 2014; Wehner et al., 2014; Brandão et al., 2019). While the importance of metabolic reprogramming in regeneration is increasingly recognized, its molecular links to cell fate determination remain poorly understood.

Epigenetic modifications, such as DNA methylation and histone post-translational modifications (PTMs), influence chromatin accessibility, thereby regulating spatiotemporal gene expression during tissue regeneration (Barrero & Izpisua Belmonte, 2011). Injury-induced gene expression relies on activation of regeneration-associated enhancers, highlighting the importance of the mechanisms underlying gene activation post-injury in tissue regeneration (Goldman & Poss, 2020). In zebrafish, DNA and histone methylation levels decrease during cellular dedifferentiation and proliferation, such as during the blastema formation in caudal fin regeneration, latter increasing to silence proliferative genes as cells redifferentiate during caudal fin (Hirose et al., 2013; Takayama et al., 2014) and heart regeneration (Ben-Yair et al., 2019). Conversely, histone acetylation activates pluripotency-related genes early in fin regeneration, later decreasing to facilitate cell redifferentiation (Pfefferli et al., 2014). DNA and histone modifications often rely on metabolites derived from glucose metabolism, linking metabolic activity to gene regulatory programmes and cell fate decisions (Z. Dai et al., 2020). Recently, lactate-derived histone lysine lactylation has emerged as a novel PTM, promoting gene transcription in various processes, including inflammation (D. Zhang et al., 2019), somatic cell reprogramming (Li et al., 2020), tumor growth (Jiang et al., 2021; Yu et al., 2021;Liu et al., 2022) and embryonic development (S. K. Dai et al., 2022; Merkuri et al., 2024). Lactate also acts as signalling molecule in various biological processes, including tissue repair by promoting an anti-inflammatory macrophage phenotype during ischemic muscle regeneration in mice (Brooks, 2020; J. Zhang et al., 2020). However, the role of lactylation in regulating cell fate transitions during regeneration has not been addressed, making it crucial to understand how histone lactylation influences tissue repair and regeneration through metabolic adaptation and epigenetic modulation.

This study is the first to report histone lactylation in zebrafish regeneration, demonstrating a significant increase in global lactylation and histone H3K18Lac levels in mesenchyme and osteoblast cells during the early stages of caudal fin regeneration. Our findings suggest that this increase is functionally regulated by glycolysis and lactate production, placing lactate-derived histone lactylation as a potential orchestrator of the initial stages of regeneration.

## Results and Discussion

### Histone lactylation is increased during caudal fin regeneration

We have previously reported that, after fin amputation, cells reprogramme their metabolic profile by increasing glycolytic activity and lactate production, required for blastema formation during the initial stage of caudal fin regeneration (Brandão et al., 2022). Considering that this metabolic shift leads to lactate accumulation, we hypothesised that histone lactylation could play a role in the process. As histone lactylation had not been studied in zebrafish regeneration, to address its potential function we started by performing a Western Blot (WB) analysis on caudal fin tissue using a pan anti-Kla antibody that targets lactylated lysines (Kla) across all proteins, including histones (D. Zhang et al., 2019). Full protein extraction showed low intensity in the histone molecular weight range, suggesting lower levels of lactylated lysines in histones relative to other proteins (Fig. 1A). However, by enriching the histone fraction, we successfully detected Kla within the molecular weight interval that includes histones, confirming the presence of histone lactylation in zebrafish caudal fin tissue (Fig. 1A).

**Figure 1.**
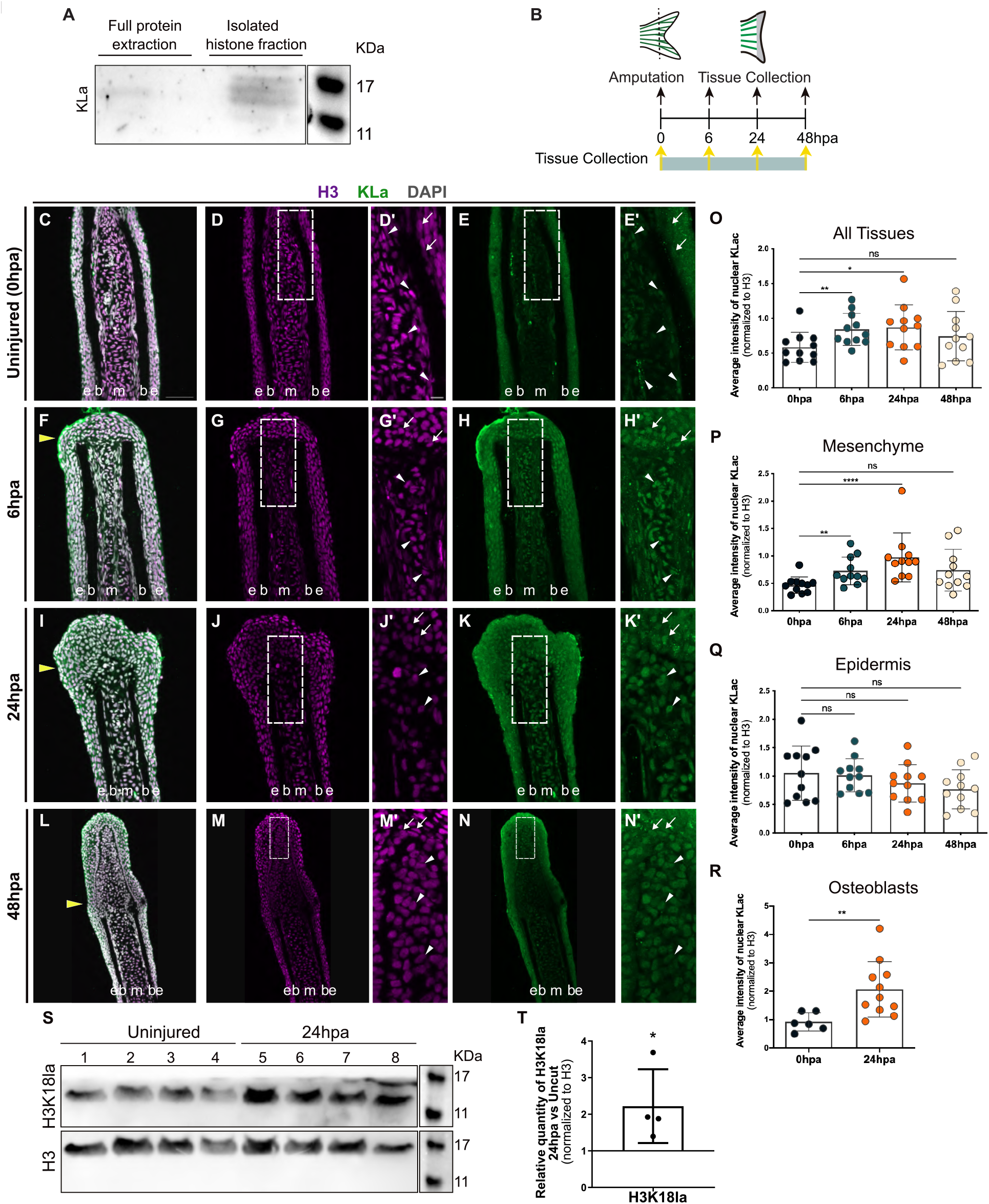
Histone lactylation increases during caudal fin regeneration. **(A)** Western blot for Kla in full protein extract and isolated histone fraction, from uninjured zebrafish caudal fin samples. Kla protein, approximately from 11 to 17 kDa in size, is detected in the corresponding histone associated region in enriched histone fraction. **(B)** Schematic representation of the caudal fin amputation assay, with the different time-points of tissue collection. **(C-N)** Representative transverse cryosections of caudal fins immunostained for H3 (magenta) and Kla (green) antibodies and counterstained with DAPI (grey), at different time points: 0 hpa (uninjured) (C-E); 6 hpa (F-H); 24 hpa (I-K); and 48 hpa (L-N). White dashed boxes delineate magnified panels in D’,E’,G’,H’,J’,K’,M’ and N’. Arrows indicate H3+Kla+ cells in the epidermis. White arrowheads indicate H3+Kla+ cells in the mesenchyme. Yellow arrowheads indicate amputation plane. e, epidermis; b, bone; m, mesenchyme. Hpa, hours post-amputation. Scale bar represents 50 μm (C) and 10 μm (D’) in magnified panels. **(O-R)** Graph showing the quantification of the average intensity of nuclear Kla, normalized to the average intensity of H3, at 0 hpa, 6 hpa, 24 hpa and 48 hpa, in the whole caudal fin tissue (O), mesenchyme (P), epidermis (Q) and osteoblasts (R). Statistical analysis corresponds to Mann-Whitney test with Mean ± SD (n=11 blastemas from 4 fish). **(S)** Western blot for H3K18la in uninjured (lanes 1-4) and 24 hpa (lanes 5-8) fins, H3 used as control. Each lane corresponds to one biological replicate. **(T)** Quantification of the relative quantity of H3K18la, normalized to H3, at 24 hpa compared to uninjured fins. Statistical analysis corresponds to Mann-Whitney test with Mean ± SD (n=4 biological replicates). ns, not significative; *P<0,05; **P<0,01; ****P<0,0001.

Considering the direct effect of lactate in lysine lactylation(D. Zhang et al., 2019), and our previous findings that lactate levels increase within the first 24 hpa (Brandão et al., 2022), we examined whether histone lactylation is altered during the initial stages of caudal fin regeneration. For this purpose, we analysed tissue lactylation levels, measuring Kla intensity by immunofluorescence at 6, 24 and 48 hpa, compared to uninjured fins (0 hpa) (Fig. 1B). Since Kla detects all lactylated proteins, we used DAPI to restrict Kla signal quantification to the nucleus (Hagihara et al., 2021). Additionally, we normalized Kla signal to the levels of histone 3 (H3) (Pfefferli et al., 2014). We observed a significant increase in nuclear lactylation intensity in the whole fin tissue at 6 and 24 hpa, compared to uninjured control (Fig. 1C-K,O), but not at 48 hpa, when lactylation levels returned to those observed in uninjured fins (Fig. 1C-E,L-O). This result suggests that histone lactylation increases during the early stages of regeneration, when cell dedifferentiation and migration are necessary for blastema formation, and then returns to basal levels by the second day of regeneration, when the blastema is fully formed. Interestingly, when analysing the main fin tissues independently, lactylation levels increase in the mesenchymal compartment at 6 and 24 hpa (Fig. 1P), compared to uninjured control, while remaining unchanged in the epidermis across all-time points analysed (Fig. 1Q). Given our previous findings that metabolic reprogramming towards glycolysis is critical for the dedifferentiation of mature osteoblast during regeneration (Brandão et al., 2022), we also examined histone lactylation in this cell type at 24 hpa, when lactylation levels in the mesenchyme are more elevated. Using o*sx*:mCherry reporter fish, which labels osteoblasts, we observed that lactylation levels also increase in these cells, compared to uninjured fins (0 hpa) (Fig. 1R, Fig. S1).

To further validate our results, we assessed the lactylation of histone H3 at lysine residue 18 (H3K18la), a modification recently identified as marking active promoters in macrophages (D. Zhang et al., 2019), cancer cells (Yu et al., 2021), reprogramming somatic cells (Li et al., 2020) and tissue-specific active enhancers (Desgeorges et al., 2024; Galle et al., 2022). We used the H3K18la specific antibody to perform a WB on uninjured (0 hpa) and 24 hpa regenerating caudal fins, showing that the H3K18la modification is present (Fig. 1S) and that the expression levels increase at 24 hpa, compared to uninjured condition (Fig. 1T), which mirrors the increase in nuclear Kla levels (Fig. 1C-K, O).

These findings highlight the importance of histone lactylation during early caudal fin regeneration, with global lactylation and histone H3K18Lac levels increasing in the first 24-hours post-amputation and returning to baseline by 48 hpa. This increase is specific to mesenchymal and osteoblast cells, suggesting that histone lactylation may reshape the epigenetic landscape, promoting cellular plasticity and enabling osteoblasts to re-enter the cell cycle, proliferate, and differentiate into new osteoblasts, thus contributing to bone formation (Brandão et al., 2019). Overall, our data suggest that histone lactylation is critical for maintaining mature osteoblast plasticity, potentially driving key regenerative processes such as cell dedifferentiation, migration, and blastema proliferation, all crucial for effective tissue regeneration.

### Histone lactylation is driven by lactate

To determine whether lactate levels directly contribute to increased histone lactylation during the first hours post-amputation (Fig. 1), we administered glucose and lactate to uninjured wild-type fish to assess whether lactylation levels could be induced, mimicking those seen in a regenerative setting. To minimize variability in lactate levels caused by normal feeding, fish were fasted for 24 hours before injection (Fig. 2A). Immunofluorescence analysis of Kla at 24 hours post-injection revealed no significant difference in the average nuclear Kla intensity levels between glucose-injected and control PBS-injected fish (Fig. 2B-G’,K). However, fish injected with lactate showed a significant increase in nuclear Kla levels, compared to controls (Fig. 2B-D’,H-K), confirming that histone lactylation responds to changes in lactate levels.

**Figure 2.**
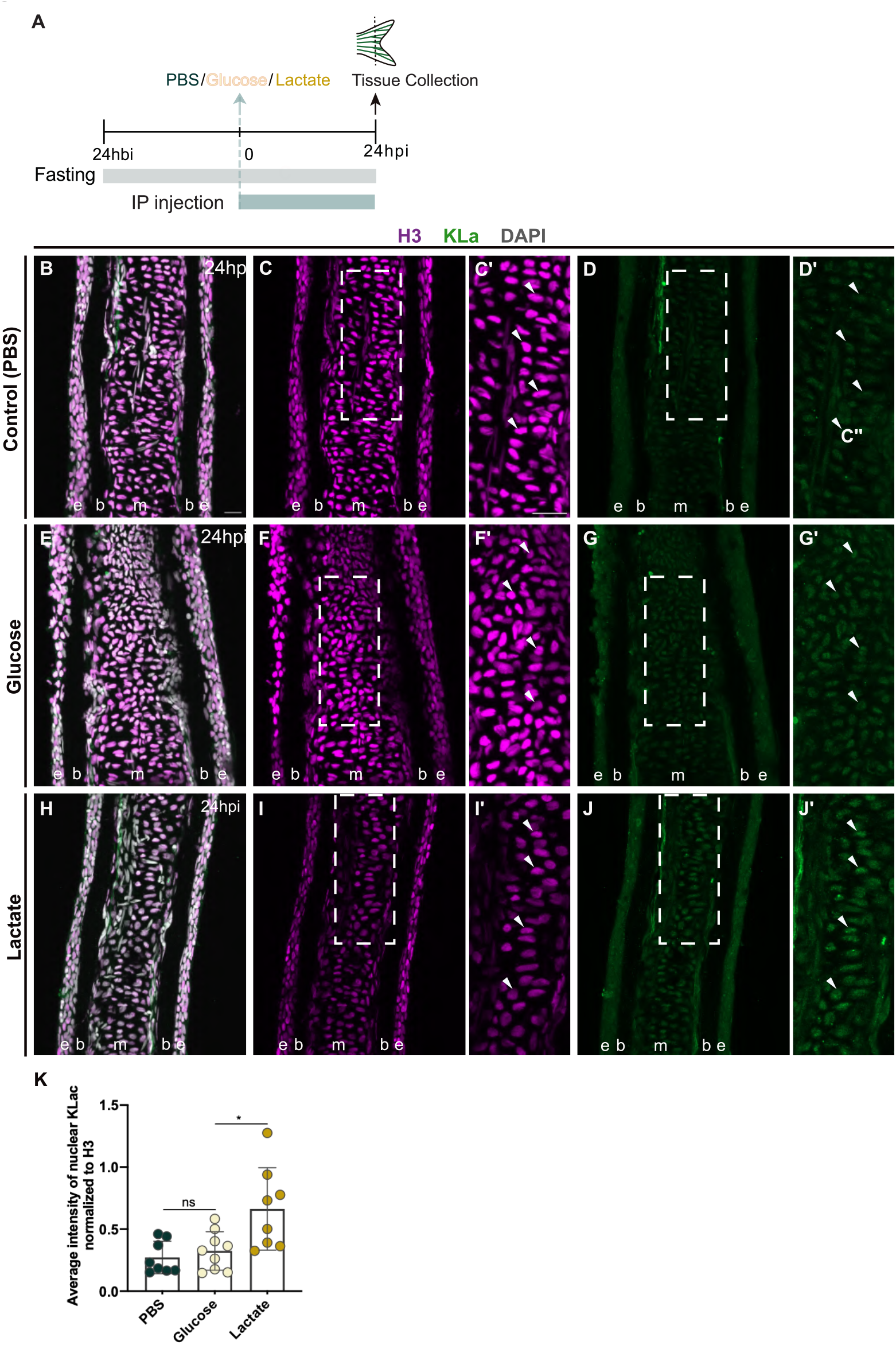
Histone lactylation is driven by lactate. **(A)** Schematic representation of the experimental design used to administer glucose and lactate. Fish were placed in a fasting condition 24 hbi, administered via IP injection with glucose and lactate and kept in fasting condition until 24 hpi, upon which the caudal fins were collected. **(B-J)** Representative transverse cryosections of 24 hpi caudal fins immunostained for H3 (magenta) and Kla (green) antibodies and counterstained with DAPI (grey), in fish treated with: PBS (control) (B-D); glucose (E-G); and lactate (H-J). White dashed boxes delineate magnified panels in C’,D’,F’,G’,I’,J’. White arrowheads indicate H3+Kla+ cells. e, epidermis; b, bone; m, mesenchyme. hours before injections (hbi); hours post-injection (hpi). Scale bar represents 100 μm (B) and 20 μm (C’) in magnified panels. **(K)** Quantification of the average signal intensity of nuclear Kla, normalized to the H3 signal. Statistical analysis corresponds to Mann-Whitney test with Mean ± SD (for control PBS and lactate, n= 8 blastemas from 3 fish; for glucose, n=9 blastemas from 3 fish). n.s., not significative; *P<0,05.

These results provide evidence that exogenous lactate can stimulate lactylation. By administering lactate to uninjured zebrafish, which naturally have lower lactate levels in the fin than in regenerating conditions (Brandão et al., 2022), we induced an increase in Kla levels (Fig. 2B,D,E), similar to what is observed during regeneration (Fig. 1O). This suggests that higher levels of histone lactylation during regeneration likely stems from elevated lactate levels associated with metabolic reprogramming. In contrast, exogenous glucose did not affect Kla levels (Fig. 2B-G’), as in the uninjured condition where no metabolic reprogramming is triggered, suggesting that glucose is mainly diverted towards oxidative phosphorylation rather than glycolysis, making it insufficient to increase lactate production.

Overall, these findings suggest that the early increase in histone lactylation during regeneration is likely driven by elevated lactate levels, consistent with findings reported in other biological systems (Hu et al., 2024; Wei et al., 2023; D. Zhang et al., 2019).

### Glycolysis and lactate generation inhibition impair histone lactylation

To further investigate the functional relevance of lactate in histone lactylation, we inhibited glycolysis and assessed its impact on histone lactylation levels following injury. We used an established glycolysis inhibition protocol (Brandão et al., 2022) (Fig. 3A), administering the glucose analogue 2-Deoxy-D-glucose (2DG), which competes with glucose for the binding of the hexokinase catalytic domain, the first enzyme in the glycolysis pathway. We treated *osx*:mCherry reporter fish with 2DG from 0 hpa until 24 hpa, period of significant increase in histone lactylation levels (Fig. 1O). Immunofluorescence analysis for Kla revealed a major decrease in nuclear Kla intensity across the entire fin tissue adjacent to the amputation plane in 2DG-treated fins, compared to controls (Fig. 3B-J). When analysing each major cell population individually, this reduction was observed in the epidermis, mesenchyme, and osteoblast populations (Fig. 3K-M).

**Figure 3.**
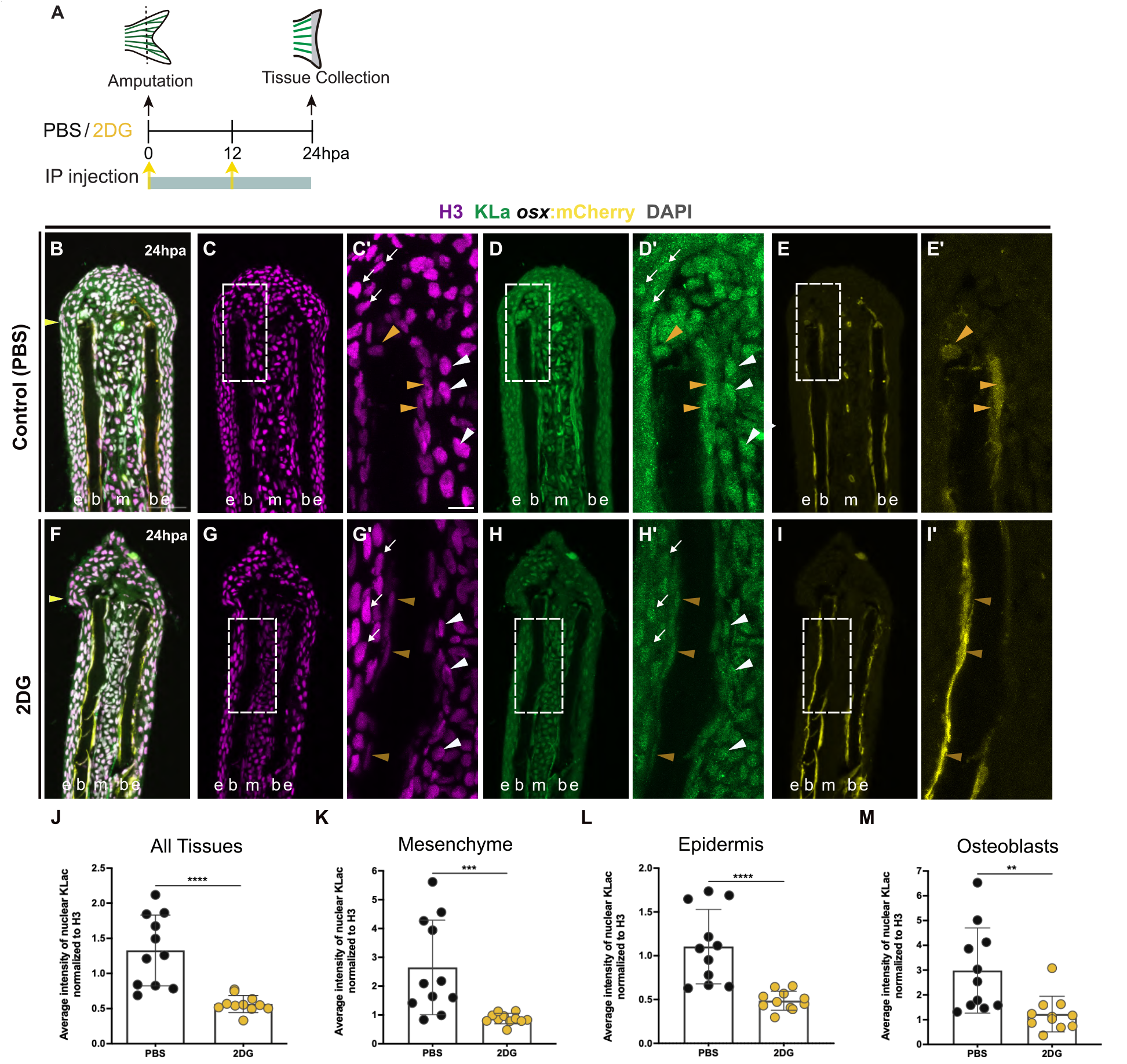
Glycolysis inhibition impairs histone lactylation. **(A)** Schematic representation of the experimental design used to inhibit glycolysis. Fish were administered, via IP injection, with vehicle (PBS) or glycolytic inhibitor, 2DG, every 12 hr, from fin amputation (0 hpa) until 24 hpa. **(B-I)** Representative transverse cryosection of 24 hpa *osx*:mCherry (yellow) caudal fins immunostained for Kla (green) and H3 (magenta) antibodies and counterstained with DAPI (grey), in fish treated with: PBS (control) (B-E) and 2DG (F-I). White dashed boxes delineate magnified panels in C’-E’, G’-I’. Arrows indicate H3+Kla+ cells in the epidermis. White arrowheads indicate H3+Kla+ cells in the mesenchyme. Orange arrowheads indicate H3+Kla+osx+ osteoblasts. Yellow arrowheads indicate amputation plane. e, epidermis; b, bone; m, mesenchyme. hpa, hours post-amputation. Scale bar represents 50 μm (B) and 10 μm (C’) in magnified panels. **(J-M)** Quantification of the average signal intensity of nuclear Kla, normalized to the H3 signal, in 2DG treated fins compared to control in the whole tissue (J), mesenchyme (K), epidermis (L) and OBs (M). Statistical analysis corresponds to Mann-Whitney test with Mean ± SD (n=11 blastemas from 4 fish). **P<0,01; ***P<0,001; ****P<0,0001.

To further validate these findings, we administered galloflavin, a known inhibitor of LDHA, to block lactate production (Farabegoli et al., 2012). Since blastema formation is a prerequisite for caudal fin regeneration and considering that we previously demonstrated that lactate production is essential for blastema growth (Brandão et al., 2022), we first assessed the impact of galloflavin on blastema growth. Galloflavin was administered at two time points post-amputation, 0 hpa and 24 hpa, and the total fin regenerated area measured at 48 hpa (Fig. 4A). We observed a significant reduction in the caudal fin regenerated area, compared to control fins (Fig. 4B-D), confirming the importance of lactate formation for tissue regeneration. Subsequently, we treated *osx*:mCherry reporter fish with galloflavin and analysed Kla immunofluorescence at 24 hpa, the time point until which we observe a significant increase in histone lactylation (Fig. 1O). Consistent with the results of glycolysis inhibition, galloflavin-treated fins showed a decrease in nuclear Kla intensity throughout the entire fin tissue, compared to control fins (Fig. 4E-M). The analysis of each major cell population individually, also revealed that nuclear Kla intensity was decreased in the epidermis, mesenchyme, and osteoblast populations (Fig. 4N-P).

**Figure 4.**
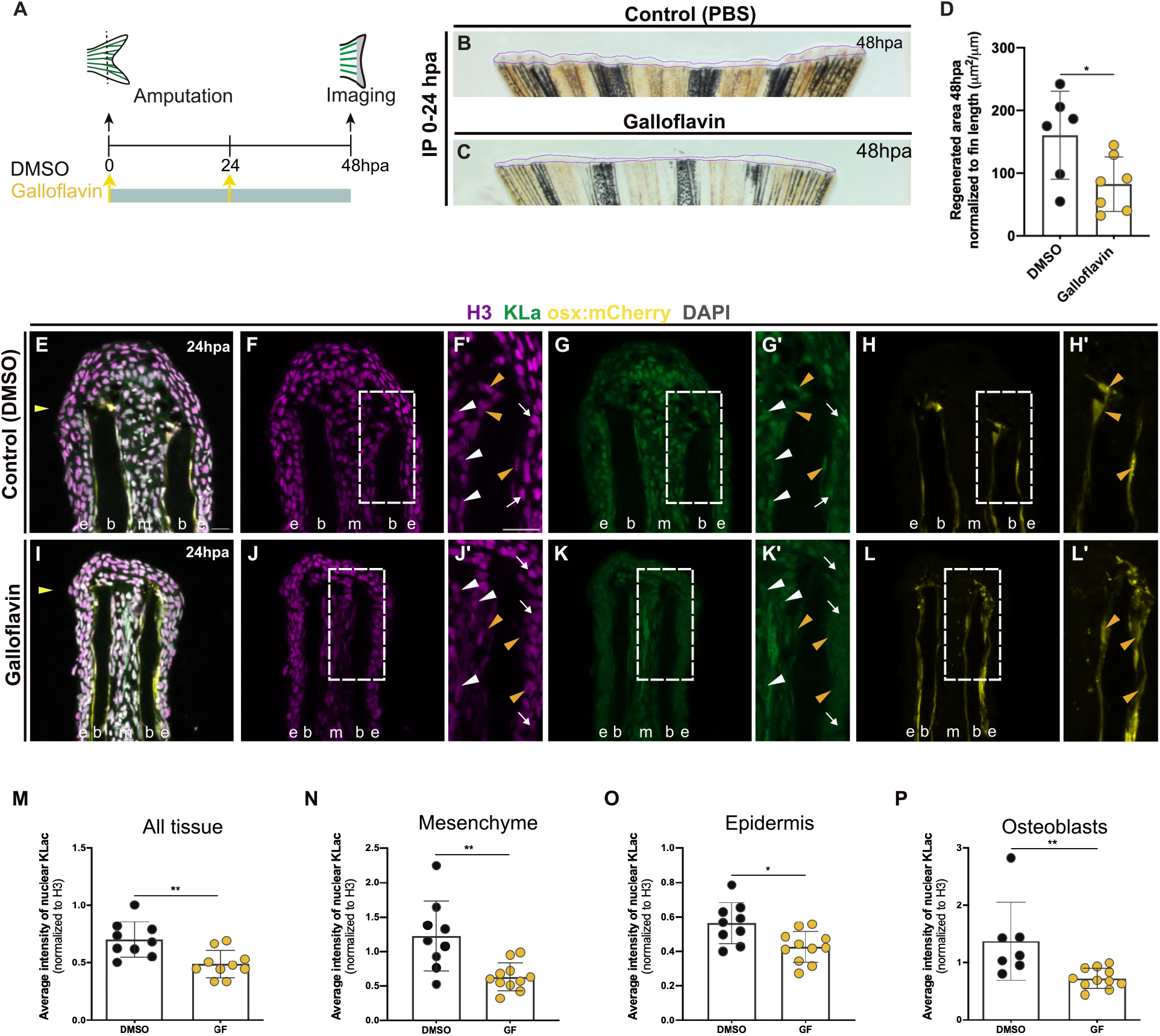
Lactate inhibition impairs histone lactylation. **(A)** Schematic representation of the experimental design used to inhibit lactate. Fish were administered, via IP injection, with vehicle (DMSO) or lactate inhibitor, Galloflavin, at fin amputation (0 hpa) and at 24 hpa. **(B**,**C)** Representative images of 48 hpa whole fins treated with vehicle (DMSO) (B) or Galloflavin (C). **(D)** Quantification of the total fin regenerated area at 48 hpa, after vehicle (DMSO) or Galloflavin injection. Statistical analysis corresponds to Mann-Whitney test with Mean ± SD (for control DMSO, n=6 fish; for Galloflavin, n=7 fish). *P<0,05. **(E-L)** Representative transverse cryosection of 24 hpa osx:mCherry (yellow) caudal fins immunostained for Kla (green) and H3 (magenta) antibodies and counterstained with DAPI (grey), in fish treated with: DMSO (control) (E-H) and Galloflavin (I-L). White dashed boxes delineate magnified panels in F’-H’, J’-L’. Arrows indicate H3+Kla+ cells in the epidermis. White arrowheads indicate H3+Kla+ cells in the mesenchyme. Orange arrowheads indicate H3+Kla+osx+ osteoblasts (orange). Yellow arrowheads indicate amputation plane. e, epidermis; b, bone; m, mesenchyme. hpa, hours post-amputation. Scale bar represents 100 μm (E) and 20 μm (F’) in magnified panels. **(M-P)** Quantification of the average signal intensity of nuclear Kla, normalized to the H3 signal, in Galloflavin treated fins compared to control in the whole tissue (M), mesenchyme (N), epidermis (O) and OBs (P). Statistical analysis corresponds to Mann-Whitney test with Mean ± SD (for control DMSO, n=9 blastemas from 3 fish; for Galloflavin, n=10 blastemas from 4 fish). *P<0,05; **P<0,01.

The decrease in histone lactylation following inhibition of glycolysis and lactate production highlights the critical role of metabolic reprogramming in regulating histone lactylation during the early stages of regeneration. These observations align with our previous findings of increased *ldha* expression and lactate production within the first 24 hpa of caudal fin regeneration, and the requirement of lactate production for blastema formation (Brandão et al., 2022). Furthermore, recent studies identify histone lactylation as a driver of immune responses to tissue injury. For instance, in macrophages, histone lactylation influences gene expression during ischemia-induced muscle damage in mice, with H3K18la marking active promoters and tissue-specific enhancers(Desgeorges et al., 2024). Similarly, lactate-induced histone lactylation in microglia promotes scar formation following spinal cord injury (Hu et al., 2024). Collectively, these findings suggest that histone lactylation plays diverse roles in the epigenetic regulation of tissue regeneration and repair, influencing both immune responses and the regeneration process itself.

In this study, we present new evidence that glycolysis regulates histone lactylation during zebrafish caudal fin regeneration, highlighting the potential role of lactylation in controlling gene expression critical for bone regeneration. Future research should explore the mechanism of lactate-derived histone lactylation, identifying its target genes, and assess their contribution to regeneration. Targeting histone lactylation may offer new therapeutic avenues for bone repair, particularly in treating injuries or conditions such as osteoporosis, by stimulating bone regeneration, remodelling, or preventing osteoblast dysfunction.

## Methods

### Ethics statement

All handling and experiments involving animals were approved by the ORBEA-NMS, Animal User and Ethical Committees at Centro de Estudos de Doenças Crónicas (CEDOC), ORBEA-Champalimaud and accredited by the Direcção Geral de Alimentação e Veterinária (DGAV) according to the directives from the EU (Directive 2010/63/UE) and National legislation (Directive 113/2013) for animal experimentation and welfare.

### Zebrafish lines maintenanceand caudal fin amputation

Wild-type (WT) AB and the transgenic zebrafish line *Tg(osterix:mCherry-NTRo)*^*pd46*^ (referred as *osx*:mCherry), kindly provided by Kenneth Poss (Singh et al., 2012), were maintained in a circulating system with a 14 hour/day and 10 hour/night cycle at 28 °C (Westerfield, 2000). All experiments were performed in 3-12 months-old fish and transgenics used as heterozygotes. Caudal fin amputations were performed in fish subjected to analgesia with 4 mg/L Lidocaine 2% Braun (Braun) and anesthetized with 160 mg/mL MS-222 (Sigma, E10521), using a sterile scalpel. Amputations were made 1 or 2 segments below the first bone-segment bifurcation, removing approximately one half of the fin, as previously described (Poss et al., 2000). Fish were left to regenerate in an incubator at 33°C with water from the circulating system and fins collected at predetermined time-points post-amputation. Regenerated fins were collected from anaesthetized fish, and either processed for protein extraction or cryosectioning.

### Protein extraction

For protein extraction, 20 caudal fins from uninjured or regenerating conditions were collected for each biological replicate and four biological replicates used per condition. Caudal fins were lysed with Tris buffer (100 mM Tris-HCl pH 9.5, 1 % SDS), homogenised through sonication using Sonifer SFX 150 – 10 cycles, 10 sec each at 30 % power, and centrifuged at 16,000 g for 10 min at 4 °C. Total protein concentration from supernatants was determined using Pierce BCA protein assay kit (Thermo Fisher Scientific). Samples were then used for Western blot (WB).

### Histone enrichment

For caudal fin samples histone enrichment protocol, to identify the profile of lactylated histones through WB, 4 pools of 60 uninjured caudal fins were collected into PBS at 4 ºC. Lysis buffer (10 mM Tris-HCl pH=6,5; 50 mM Sodium Bisulphate; 0,1 % Triton X-100, 10 mM MgCl; 8,6 % sucrose – all from Sigma-Aldrich) was added and cells homogenised by applying 20 strokes in a tight fitting Dounce homogeniser. Released nuclei were pelleted by centrifugation at 2500 g, for 10 min, at 4 ºC. Pellet was then centrifuged in lysis buffer, washed in washing buffer (10 mM Tris-HCl; 13 mM EDTA pH=7.4 – all from Sigma Aldrich), and centrifuged again at 2500 g for 10 min, each. Remaining pellet was resuspended in cold H2SO4 (Sigma-Aldrich, 0,4 M) and incubated for 1 hour at 4 ºC. For final nuclear precipitation, mixture was centrifuged and 1 mL of cold acetone added to the supernatant, followed by an overnight (ON) incubation at -20 ºC. Samples were then centrifuged, pellet air-dried and resuspended in distilled water. Nuclear protein concentration was determined using Pierce BCA protein assay kit (Thermo Fisher Scientific).

### Western blot

For each sample, 20 μg of protein was denatured in Laemmli buffer and heated at 95 °C for 5 min. Total protein was resolved on a 15 % SDS gel (for 10 mL: 2,3 mL MiliQ water; 5 mL 30 % Acrylamide/Bis Solution (37:5:1); 2,5 mL of separating buffer (90 g Tris, 2 g SDS in 500 mL water, pH= 8.8); 0,1 mL of Ammonia Persulfate 10 % and 10 μL of TEMED)) and transferred to a nitrocellulose membrane at 100 V during 75 min. Membranes were blocked with Tris buffered saline (TBS) with 0.1 % Tween 20 (TBST) containing 5 % non-fat dry milk, followed by incubation with primary antibodies (for antibody details see Table S1), ON at 4 °C. Membranes were washed in TBST and incubated with HRP-conjugated secondary antibodies for 1 hour at RT. For imaging, membranes were incubated with ECL TM Prime Western Blotting Detection Reagent (Cytiva) and images acquired using ChemiDoc Touch System (BioRad) within the linear range.

### Pharmacological and chemical treatments

For pharmacological treatments fish were subjected to intraperitoneal injections (IP) at designated time-points with either 2DG (Sigma-Aldrich, 0,5 mg/g diluted in 1x Phosphate Buffered Saline (PBS)); Galloflavin (Cayman, 0,12 mg/g diluted in DMSO); Glucose (Sigma-Aldrich, 3 mg/g); Sodium Lactate (Sigma-Aldrich, 3mg/g) or with corresponding vehicle (Control). IP injections were performed with an insulin syringe U-100 G 0,3 mL and a 30 G needle (BD Micro-fine) inserted close to the pelvic girdle. For all experiments water was replaced daily and fish left to regenerate until the desired time-point.

### Immunofluorescence and image acquisition

Tissue processing for cryosections was performed as preciously described (Brandão et al., 2022). Shortly, fins were collected, fixed overnight (ON) in 4 % paraformaldehyde (in 1x PBS) and stored in 100 % methanol (MeOH) at -20 °C, until required. They were then gradually rehydrated in a series of MeOH/1x PBS (75 %, 50 % and 25 %) and incubated ON in 30 % sucrose (Sigma-Aldrich, in 1x PBS). Subsequently, fins were embedded in 7,5 % gelatin (Sigma-Aldrich)/ 15 % sucrose in 1x PBS and subsequently frozen in isopentane at -70 ºC and stored at -80 ºC. Longitudinal caudal fins sections were obtained at 12 μm using a Microm cryostat (Cryostat Leica CM3050 S) and slides stored at -20 °C. For immunofluorescence on cryosections, slides were thawed for 15 min at room temperature (RT), washed twice in 1x PBS at 37 °C for 10 min and subjected to an antigen retrieval step consisting in a 15 min incubation at 95 ºC with heated sodium citrate buffer (10 mM Tri-sodium citrate with 0,05% Tween20, pH6). Slides were then incubated in 0.1 M glycine (Sigma-Aldrich, in 1x PBS) for 10 min, permeabilized with acetone for 7 min at -20 ºC and incubated for 20 min in 0,2% PBST (1x PBS with 0,2 % TritonX-100). Afterwards, they were incubated in a blocking solution (10 % non-fat dry milk in PBST) for at least 2hours at RT. Slides were incubated with primary antibodies diluted in blocking solution, ON at 4 ºC (for primary antibody details see Table S1). Next day slides were washed with PBST 6 times, 10 min each, and incubated with secondary antibodies diluted in blocking solution, for 2 hours at RT, in the dark (for secondary antibody details see Table S2). Slides were then washed in PBST three times, 10 min each, and counterstained with 4’,6-diamidino-2-phenylindole (DAPI; 0.001mg/mL in 1x PBS, Sigma-Aldrich) for 5 min in the dark, for nuclei staining. Slides were washed three times with 1x PBS, 10 min each, mounted with fluorescent Mounting Medium (DAKO) and stored at 4ºC protected from light until image acquisition.

### Image acquisition and processing

For regenerated area measurements, images of live anesthetised WT adult caudal fins were acquired using a Zeiss Discovery.V8 stereoscope equipped with a Zeiss AxioCam ICc1 camera using an objective Achromat S 1.0x FWD 63 mm (at 1x zoom) controlled by Zen 2 PRO blue software. Images were acquired using transmitted light filters and assembled using the Fiji software (Schindelin et al., 2012).

Immunolabeled cryosections were analysed in confocal microscope Zeiss LSM 980 controlled by ZEN 3.3. Cryosection images were acquired using a C-Apochromat 40×1.2 NA water objective with 0.6 x zoom, a step size of 1 μm, and 488, 568, 633 and 647 nm excitation wavelengths coupled with transmitted light PMT. Sequential images were acquired to capture the first segment below the amputation plane and the entire regenerated region. For image analysis and processing, composite maximum intensity z-stack projections were made using the Fiji software (Schindelin et al., 2012). When required, concatenation of several images along the proximal-distal axis of the same longitudinal section was performed using the Fiji plugin 3D Pairwise Stitching (Schindelin et al., 2012).

For all cryosections of manipulated fish and corresponding control, images were acquired employing identical settings (magnification, contrast, gain and exposure time) and in identical/comparable regions. All Images were then processed using the Inkscape (open source <u>vector graphics editor</u>)

### Quantification and statistical analysis

For WB quantifications, 4 biological replicates were analysed per condition. Intensity of each band was quantified by densitometry using the Analyse gels tool on FIJI software. Statistical significance between the different time-points post-amputation was determined by using the Mann-Whitney U test, a non-paired, non-parametric comparison, in the Prism Graphpad software, version 9. Only p-values < 0,05 were considered statistically significant.

Measurements of total regenerated fin area were obtained by delineating distal end of the regenerated area using the Area tool in Fiji. These were normalized to the total caudal fin width, to avoid discrepancies between different fish sizes, thus resulting in one measurement value per animal.

To determine average intensity of KLac and H3 in complete longitudinal cryosections of individual regenerating bony-rays, we used IMARIS software (version 10.2.0). We used the surface tool for DAPI channel, with a nucleus surface diameter detail of 0,5 μm. This gives us the medium intensity values of all channels colocalizing with the nucleus, including KLac and H3. To independently address nuclear KLac and H3 average intensity values of the epidermis and mesenchyme, we used the surface tool, with manual drawing option to delineate the desired area (either the epidermis or mesenchyme), and then used the mask tool to create a duplicated mask of selected area. We then used the surface tool for DAPI channel on the duplicated mask of selected area, with a nucleus surface diameter detail of 0,5 μm. This gives us the medium intensity values of all channels colocalizing with the nucleus, including KLac and H3, in the duplicated selected area. To address nuclear KLac and H3 average intensity values in osteoblasts, we used surface tool for mCherry channel, with a surface detail of 0,5 μm and then used the mask tool to create a duplicated mask of selected osx:mCherry area. We then used the surface tool for DAPI channel on the duplicated mask of osx:mCherry area, with a nucleus surface detail of 0,5 μm. This gives us the medium intensity values of all channels colocalizing with the nucleus, including KLac and H3, in the duplicated osx:mCherry selected area. For each experiment the average intensity of KLac was normalized for the average intensity of H3. For each quantification, 3-6 animals were used per condition and cryosections corresponding to at least 3 bony-rays per animal were analysed. The exact number of animals and cryosections used is discriminated in the corresponding figure legend. Mean ± SD are displayed in the graphs.

For all adult caudal fin regenerate measurements and quantifications performed on cryosections, statistical significance between controls and manipulated animals, or between different time-points post-amputation or time-intervals, was determined by using the Mann-Whitney U test, a non-paired, non-parametric comparison, in the Prism Graphpad software, version 6 and 9. Only p-values < 0,05 were considered statistically significant.

## Acknowledgements

We are grateful to Ana Teresa and Lara Carvalho for advice and for reading the manuscript, Telmo Pereira for data analysis and Fior Lab from CF for providing laboratory space and materials. We thank Petra Pintado and Fábio Valério from the NMS Fish Facility, and Catarina Certal, Joana Monteiro, et al., from the CF Fish Platform for animal care; Ana Farinho from the NMS Histology facility, and Sérgio Casimiro, Inês Romano and Ana Quitéria from the CF Histopathology Platform for assistance in tissue processing and cryosectioning; and Telmo Pereira from NMS Microscopy Facility for technical assistance.

## Author contributions

Conceptualization, J.B., R.L. and A.S.B.; Methodology, J.B., R.L., A.S.C. and A.S.B.; Investigation, J.B. and R.L.; Writing – Original Draft, R.L.; Writing – Review & Editing, R.L., J.B. A.S.C., R.M., and A.J.; Funding Acquisition, A.J.; Resources, RM and A.J.; Supervision, A.S.B., R.L and A.J.

## Data and resource availability

All relevant data and resource can be found within the article and its supplementary information.

## Supplemental information

Document S1. Figure S1, Table S1 and S2

