## Supplementary material for "Metabolic reprogramming regulates histone lactylation during zebrafish caudal fin regeneration": Figure S1

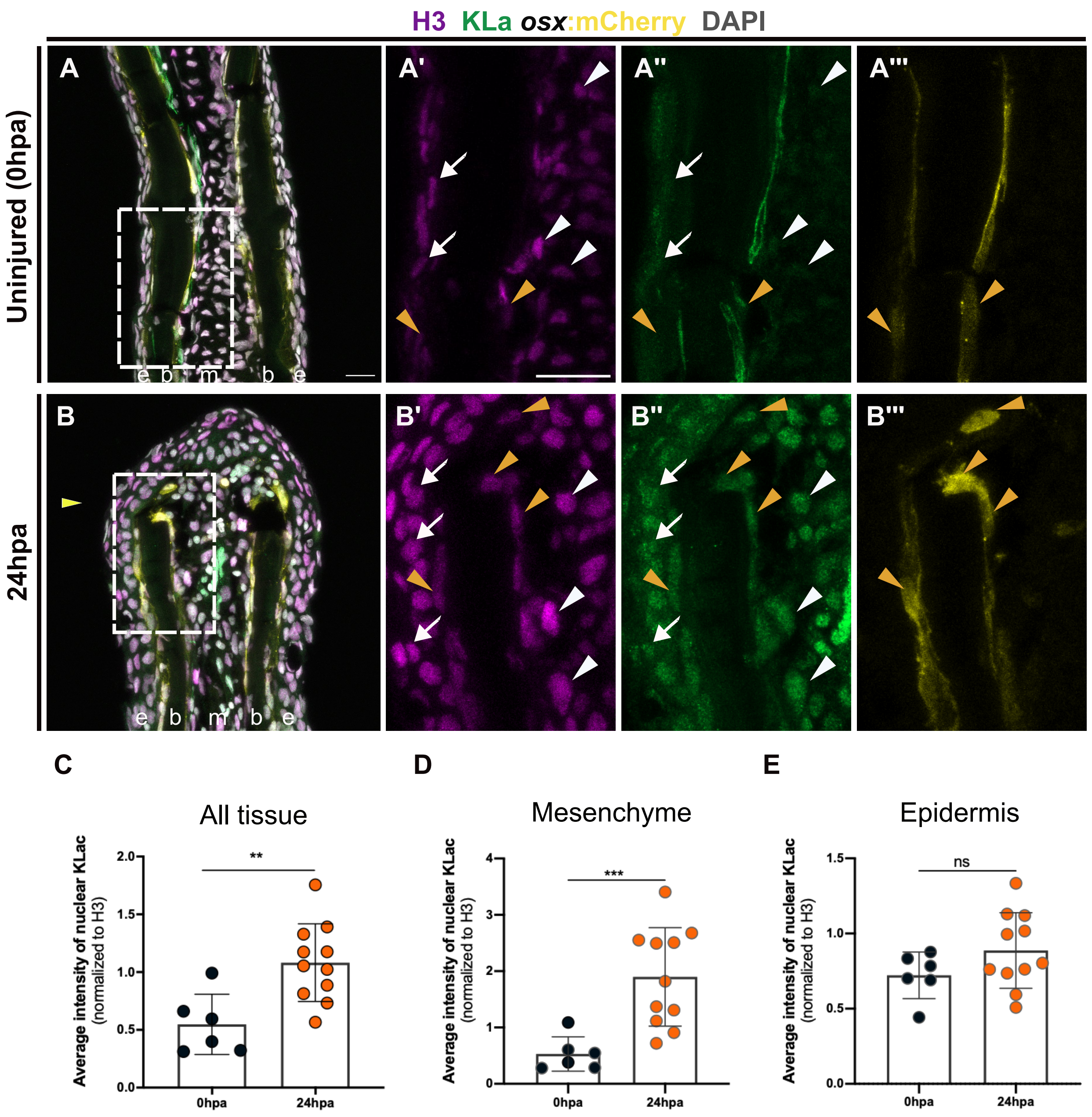


**Figure S1. Increase of histone lactylation in osteoblasts during caudal fin regeneration.** (A,B) Representative transverse cryosections of osx:mCherry (yellow) caudal fins immunostained for Kla (green) and H3 (magenta) antibodies and counterstained with DAPI (grey), at different time points: 0 hpa (uninjured condition) (A) and 24 hpa (B). White dashed boxes delineate magnified panels in A'-A''', B'-B'''. Arrows indicate H3+Kla+ cells in the epidermis. White arrowheads indicate H3+Kla+ cells in the mesenchyme. Orange arrowheads indicate H3+Kla+osx+ osteoblasts. Yellow arrowheads indicate amputation plane. e, epidermis; b, bone; m, mesenchyme. Hpa, hours post-amputation. Scale bar represents 100 µm (A) and 20 µm (A') in magnified panels. (C-E) Graph showing the quantification of the average intensity of nuclear Kla, normalized to the average intensity of H3, at 0 hpa and 24 hpa in the whole caudal fin tissue (C), mesenchyme (D) and epidermis (E). Statistical analysis corresponds to Mann-Whitney test with Mean ± SD (for uninjured, n=6 blastemas from 3 fish; for 24 hpa, n=11 blastemas from 5 fish).

**Table S1:** List of primary antibodies used.

| Antibody | Host | Dilution | Assay | Company |
|---|---|---|---|---|
| Pan anti-Kla | Rabbit | 1:200 | IF/WB | PTM-BIO, PTM-1401 |
| Anti-Histone H3 | Mouse | 1:200 | IF | Santa Cruz Biotechnology, SC-517576 |
| Anti-mCherry | Rat | 1:200 | IF | Thermo Fisher Scientific, M11217 |
| Anti-Histone H3 | Rabbit | 1:500 | WB | Milipore, 06-755 |
| Anti-H3K18la | Rabbit | 1:500 | WB | PTM-BIO, PTM-1406 |

**Table S2:** List of secondary antibodies used.

| Antibody | Host | Specificity | Dilution | Assay | Company |
|---|---|---|---|---|---|
| Alexa Fluor 488 | Goat | Rabbit | 1:500 | IF | Invitrogen, A11070 |
| Alexa Fluor 568 | Goat | Mouse | 1:500 | IF | Invitrogen, A11031 |
| Alexa Fluor 568 | Goat | Rat | 1:500 | IF | Invitrogen, A11077 |
| Alexa Fluor 647 | Donkey | Mouse | 1:500 | IF | Jackson Immunoresearch, 715-605-151 |
| Horseradish peroxidase-conjugated | Goat | Rabbit | 1:5000 | WB | Bio-Rad, 170-6515 |
| Horseradish peroxidase-conjugated | Goat | Mouse | 1:5000 | WB | Bio-Rad, 170- 6516 |
